# Potent and selective ‘genetic zipper’ method for plant protection: innovative DNA psyllidicides against *Trioza alacris* Flor based on short unmodified antisense oligonucleotides targeting rRNA of the pest

**DOI:** 10.1101/2024.06.02.597012

**Authors:** V.V. Oberemok, I.A. Novikov, E.V. Yatskova, A.I. Bilyk, A.K. Sharmagiy, N.V. Gal’chinsky

## Abstract

Chemical insecticides increased the chemical burden on natural ecosystems posing environmental health risk factor. The urgent need for a more sustainable and ecological approach has produced many innovative ideas, including eco-friendly ‘genetic zipper’ method (or CUAD platform) based on contact oligonucleotide insecticides. Oligonucleotide insecticides have enjoyed success recently on many sternorrhynchans showing highly adaptable structure for distinct insect pest species and selective mode of action. In this article, we describe the efficiency of the oligonucleotide insecticides (briefly, olinscides or DNA insecticides) Alacris-11 and Laura-11, as well as their combined use in mixture (1:1), designed for control of bay sucker (*Trioza alacris* Flor), an important psyllid pest of noble laurel (*Laurus nobilis* L.). These olinscides are based on short unmodified antisense DNA oligonucleotides that target ITS2 between 5.8S rRNA and 28S rRNA in pre-rRNA (Laura-11) and 28S rRNA region in mature 28S rRNA and pre-rRNA (Alacris-11). The maximum pest mortality, observed on 14^th^ day of the experiment, comprised 95.01 ± 4.42 % for Alacris-11, 97.16 ± 2.48 % for Laura-11, and 98.72 ± 1.14 % for their mixture (1:1). The control oligonucleotide CTGA-11 did not cause any significant mortality (9.38 ± 0.57 %), emphasizing selectivity in the action of oligonucleotide insecticides. The results show potent and specific nature of oligonucleotide insecticides for pest control and opens up new frontiers in control of economically important psyllids in agriculture and forestry, including Asian citrus psyllid (*Diaphorina citri* Kuwayama) and many others. Scientists can easily adopt ‘genetic zipper’ method for plethora of insect pests because DNA is a programmable molecule and provides game-changing characteristics for plant projection.

## Introduction

It is widely accepted that without chemical pesticides and host-plant resistance (including genetically-modified crops), crop losses would reach levels of 50–80% in many instances (Oerke, 2006). At the same time, hidden costs of side effects of use of chemical pesticides difficult to enumerate in economic terms, while costs connected with implementation of eco-friendly pesticides are more visible and require not only huge expenses, but also a restructuring of thinking and agricultural habits. Externally applied pesticides remain the cornerstone of pest management in most food production systems (Isman, 2019). Pesticide resistance problem pushes researchers to discover new pesticides, particularly insecticides. In our opinion, this fact should be perceived as a constant opportunity to create new insecticides with eco-friendlier characteristics and gradually replace toxic chemical insecticides out of the market.

Current environmental risks and insecticide resistance problem have generated innovative ideas among scientists in plant protection, one of them is eco-friendly ‘genetic zipper’ method based on the use of short contact unmodified antisense DNA (CUAD) targeting mature rRNA and/or pre-rRNA of insect pests (Oberemok et al. 2024). Short antisense DNA serves as contact insecticide that penetrates insect integument, enters cells and targets target rRNA in nucleus, precisely nucleolus, where ribosome biogenesis takes place. A target rRNA and/or pre-rRNA and an olinscide interlock and resemble zipper mechanism performed by DNA-RNA duplex (Figure 1). This ‘genetic zipper’ method originated in 2008 on lepidopterans (Oberemok, 2008), was rethought and expanded in 2019 (Oberemok et al. 2019) and today shows amazing results on sternorrhynchans (scale insects, mealybugs, aphids, etc.) (Oberemok et al. 2020; Oberemok et al. 2022; Puzanova et al. 2023; Gal’chinsky et al. 2024). Olinscides act through DNA containment mechanism (‘arrest’ of target rRNA, block of functioning and biogenesis of ribosomes accompanied with hypercompensation of expression of target rRNA and subsequent degradation of target rRNA by RNase H) (Gal’chinsky et al. 2024; Oberemok and Gal’chinsky 2024; Oberemok et al. 2024). DNA containment (DNAc) mechanism is supposed to play a fundamental natural role in antiviral defense by targeting the invading viral transcripts of DNA viruses for cleavage and also can be recruited by DNA viruses themselves to regulate biogenesis of rRNA (hypercompensation of rRNA during 1st step of DNAc) (Gal’chinsky et al. 2024; Oberemok et al. 2024; Oberemok, 2024). Use of insect pest pre-rRNA and mature rRNA as target leads to high efficiency of oligonucleotide insecticides, since pre-rRNA and mature rRNA comprise 80% of all RNA in the cell (Warner, 1999; Alberts et al. 2002). Thousands of different mRNAs make up only 5% of all RNA and use of mature rRNA and pre-rRNA for targeting substantially increases signal-to-noise ratio, ca. 10^5^:1 (rRNA vs. random mRNA) (Palazzo and Lee 2015). In addition to the high biodegradability potential, low carbon footprint, and safety for non-target organisms, the opportunity of solving the problem of target-site resistance using oligonucleotide insecticides based on conservative gene sequences is expected (Oberemok et al. 2023). Insect pests undergo microevolution and constantly adapt to various environmental factors, including chemical insecticides. On the contrary, none of the current classes of chemical insecticides is able to quickly adapt to the rapidly emerging resistance of insects, that is, there is no algorithm for the rapid design of new chemical insecticides. Basically, the options are sorted through trial-and-error method. The use of antisense DNA sequences shows hope in this regard.

**Figure 1.**
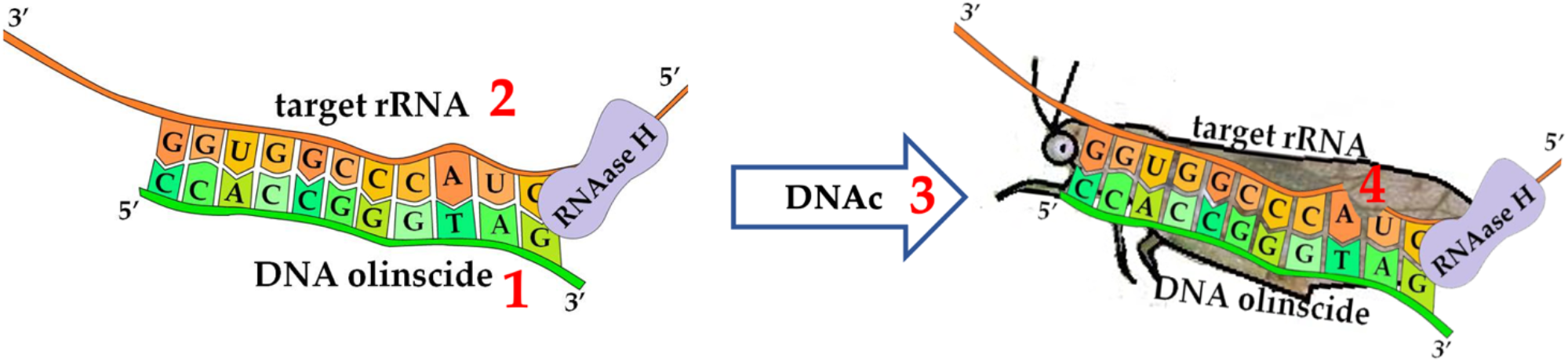
Key features of ‘genetic zipper’ method (DNA-RNA duplex formed by olinscide Alacris-11 and target sequence of pre-rRNA of *Trioza alacris)*: 1 – short unmodified antisense DNA as insecticide; 2 – rRNA as a target molecule; 3 – DNA containment (DNAc) as mechanism of action; 4 – species-specific insecticidal effect

In this article, for the first time we will use mixture of oligonucleotide insecticides against psyllids using the bay sucker *Trioza alacris* Flor as a model insect pest. Bay sucker is a psyllid from the suborder Sternorrhyncha (order Hemiptera). It feeds on economically valuable plants from the family Lauraceae, including noble laurel (*Laurus nobilis*), Azorean laurel (*L. azoria* (Seub.) Franco), Canarian laurel (*L. novocanariensis* Rivas Mart., Lousã, Fern. Prieto, E. Días, J.C. Costa and C. Aguiar), camphor (*Cinnamomum camphora* (L.) J. Presl), and Indian persea (*L. indica* L.) (Psyl’list, 2022). Among the laurel trees, only *L. nobilis* grows in Crimea, where it is one of the oldest cultivated plants. In addition to its industrial value (Zeković et al. 2009), it has great decorative value and is used for landscaping gardens, parks, public gardens, personal plots, and in hedges or in single plantings (Paparella et al. 2022). *T. alacris* causes significant damage to laurel, attacking young leaves and shoots and forming false galls on them. One gall can contain more than 15 larvae and nymphs of different ages, feeding on the sap of the plants. Fresh galls are pale green with a pink or red tint (Figure 2). Damaged leaves turn black and dry out, and the plant loses its decorative appearance.

**Figure 2.**
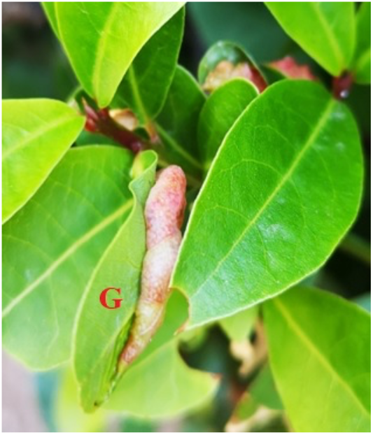
Gall (G) on the noble laurel leaf caused by *T. alacris*

**Figure 2.**
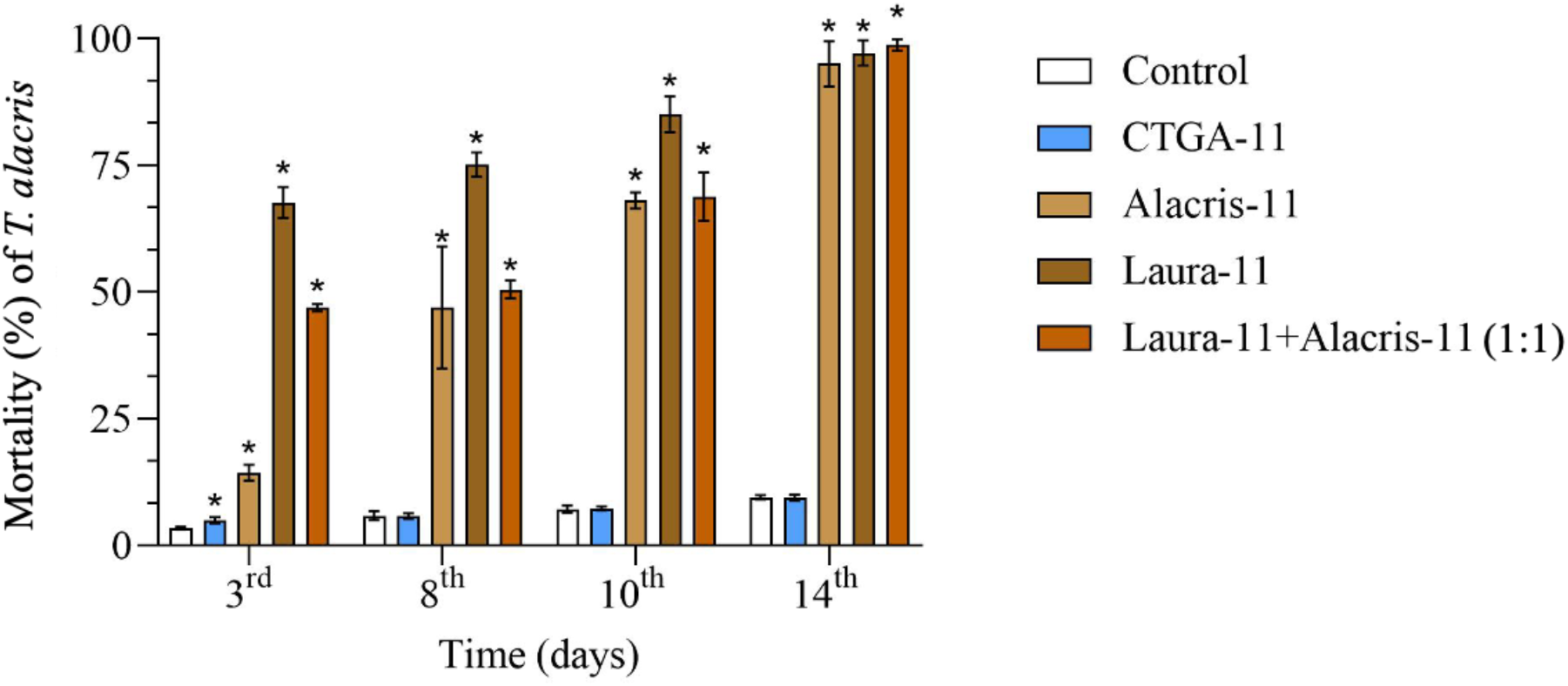
Dynamics of *T. alacris* mortality following treatment with oligonucleotide insecticides

## Materials and methods

### Sequences of oligonucleotide insecticides

Two target antisense oligonucleotides, Laura-11 (5’-GACACGCGCGC-3’) and Alacris-11 (5’-CCACCGGGTAG-3’), were designed to target the intergenic non-transcribed spacer (ITS2) between 5.8S and 28S ribosomal RNA (Laura-11) and 28S rRNA region in pre-rRNA and mature 28S rRNA (Alacris-11), respectively. GenBank (Sequence ID: MT038972.1; range: 127 to 117, Laura-11; range: 546 to 536, Alacris-11) was used for design of olinscides. Oligonucleotide CTGA-11 (5’-CTGACTGACTG-3’) was used as random sequence.

### Synthesis of oligonucleotides

Oligonucleotides were synthesized on automatic DNA synthesizer ASM-800E (BIOSSET, Novosibirsk, Russia) using standard phosphoramidite chemistry on the universal solid support UnyLinker 500Å (ChemGenes, Wilmington, USA). Cleavage and deprotection were carried out at 55 ºC overnight using a concentrated solution of ammonia. The solution was then filtered and evaporated on a vacuum rotary evaporator (Heidolph Instruments GmbH & Co. KG, Schwabach, Germany). The resulting solid substance was dissolved in deionized water (Millipore, Molsheim, France) to the required concentration, which was determined by measuring it on a spectrophotometer NanoDrop Light (Thermo Fisher Scientific, Waltham, USA).

The mass of the synthesized DNA sequences was determined using mass spectrometry MALDI-TOF analysis. Oligonucleotides were measured as positive ions with 3-hydroxypicolinic acid as the matrix on a LaserToF LT2 Plus mass spectrometer (Scientific Analysis Instruments, Manchester, UK) in a 2:1 ratio. The theoretical m/z ratio was calculated using the ChemDraw 18.0 program (ChemDraw, CambridgeSoft, USA).

Measurement of the correspondence of the synthesized oligonucleotides, determined using the MALDI-TOF method, showed that all oligonucleotides corresponded to their structure, and the resulting m/z ratio differed from the theoretically calculated m/z ratio by no more than 10 units (Table 1).

**Table 1.** Results of analysis of synthesized oligonucleotides using MALDI-TOF method.

| Oligonucleotides | Theoretical m/z ratio | Received m/z ratio |
| --- | --- | --- |
| Alacris-11 | 3341.60 | 3337.28 |
| Laura-11 | 3326.60 | 3328.73 |
| CTGA-11 | 3331.60 | 3332.48 |
| Primer Alacris-F | 4896.25 | 4898.64 |
| Primer Alacris-R | 6089.07 | 6081.59 |

### Application

Field treatments were carried out on the southern coast of Crimea. *T. alacris* larvae were treated using three oligonucleotides: the target antisense oligonucleotides Alacris-11 and Laura-11 separately and in 1:1 of their mixture; oligonucleotide CTGA-11 was used as a control sequence. The treatment was carried out on noble laurel plants using a hand-held sprayer with solution of oligonucleotides in water (100 mg/L). Experiment was performed in triplicate. The effect of treatment with olinscides was recorded between 3^rd^ and 14^th^ day of the experiment. Insect mortality was determined by calculating the ratio of dead insects to the total number of insects per 10 leaves.

### PCR

For PCR analysis, total RNA was isolated from bay sucker larvae after treatment with oligonucleotides for each recording day of the experiment using the ExtractRNA reagent (Evrogen, Moscow, Russia). For cDNA synthesis, we used a reverse transcription kit (Syntol, Moscow, Russia) according to manufacturer’s instructions. Real-time PCR was performed to assess the concentration of target rRNA using SYBR Green Master Mix kit (Roche, Basel, Switzerland) in accordance with the manufacturer’s recommendations on LightCycler® 96 Real-Time PCR System (Roche, Basel, Switzerland). Two primers were used in the PCR reaction for assessment of target rDNA expression: forward Alacris-F (5’-GAC CTC GGG CTG TAC G-3’) and reverse Alacris-R (5’-CGC TTA TTA ATA TGC TTA AA-3’). The primer annealing temperature comprised 52 °C. PCR was performed in triplicate.

### Statistical analysis

For statistical analysis, the standard error of the mean was determined and assessed using Student’s t-test. To assess the significance of differences between means of the groups also non-parametric Pearson chi-square test (χ^2^) with Yates’s correction (GraphPad Software Inc., Boston, USA) was used.

## Results

### Mortality of T. alacris after treatment with olinscides

On the 3^rd^ day of the experiment, significant insect pest mortality, 67.61 ± 3.09 %, and 14.28 ± 1.61 %, was observed after treatment of the noble laurel plants with olinscides Laura-11 and Alacris-11, respectively (p < 0.05). We also observed significant insect pest mortality, 46.86 ± 0.74 %, on the 3^rd^ day when 1:1 mixture of the olinscides (Alacris-11 and Laura-11) was used (p < 0.05). The maximum mortality of bay sucker larvae was observed on the 14^th^ day of the experiment. Mortality comprised 95.01 ± 4.42 % for group with olinscide Alacris-11, 97.16 ± 2.49 % for group with olinscide Laura-11, and 98.72 ± 1.14 % for group with their 1:1 mixture (p < 0.05). Of note, olinscide Laura-11 showed quicker dynamics in insecticidal effect than Alacris-11. As a hypothesis, we think this could be due to the fact that Laura-11 can target both, pre-rRNA and mature rRNA containing 28S rRNA, while Alacris-11 can target only ITS2 in pre-rRNA. As expected, random oligonucleotide CTGA-11 did not show any significant insecticidal effect throughout the experiment. Mortality from the application of random oligonucleotide CTGA-11 corresponded to the normal mortality in the Control group (Figure 2) and comprised 9.39 ± 0.57 % and 9.45 ± 0.41 % in CTGA-11 group and water-treated Control on the 14^th^ day of the experiment, respectively (p > 0.05).

### Target rDNA expression of T. alacris

During the experiment, expression of target rRNA was investigated. On the 3^rd^ day of the experiment, hypercompensation of target rRNA was observed for all olinscides applied (1^st^ step of DNAc mechanism) (Gal’chinsky et al. 2024). Compared to the control, expression of target rRNA increased by 3.56 ± 0.66 and 2.93 ± 0.16 for oligonucleotide olinscide Laura-11 and olinscide Alacris-11, respectively; and was 5.52 ± 0.13 times higher compared to water-treated Control for 1:1 of their mixture (Table 2).

**Table 2.** Relative concentration of target rRNA in *T. alacris* larvae after treatment with DNA oligonucleotides (Control is taken as 1).

| Day of the experiment | Target rRNA concentration |  |  |  |  |  |
| --- | --- | --- | --- | --- | --- | --- |
|  | Laura-11 |  | Alacris-11 |  | Laura-11 + Alacris-11 (1:1) |  |
| 3<sup>rd</sup> | $3.56 \pm 0.66$ | 1<sup>st</sup> step of DNAc | $2.93 \pm 0.16$ | 1<sup>st</sup> step of DNAc | $5.52 \pm 0.13$ | 1<sup>st</sup> step of DNAc |
| 8<sup>th</sup> | $0.21 \pm 0.01$ | 2<sup>nd</sup> step of DNAc | $1.35 \pm 0.01$ | 2<sup>nd</sup> step of DNAc | $0.64 \pm 0.02$ | 2<sup>nd</sup> step of DNAc |
| 10<sup>th</sup> | $0.004 \pm 0.0001$ | 2<sup>nd</sup> step of DNAc | $0.92 \pm 0.03$ | 2<sup>nd</sup> step of DNAc | $0.01 \pm 0.0004$ | 2<sup>nd</sup> step of DNAc |
| 14<sup>th</sup> | $0.58 \pm 0.05$ | 1<sup>st</sup> step of the next cycle of DNAc | $7.19 \pm 0.19$ | 1<sup>st</sup> step of the next cycle of DNAc | $0.0003 \pm 0.00002$ | 2<sup>nd</sup> step of DNAc |

On the 8^th^ and 10^th^ day of the experiment, a pattern characteristic for action of RNase H-recruiting antisense oligonucleotide was observed with a decrease in expression of target rRNA in comparison with the 3^rd^ day (2^nd^ step of DNAc mechanism). On the 14^th^ day expression of target rRNA began to increase in Laura-11 and Alacris-11 groups, while in the group of their 1:1 mixture target rRNA expression was significantly down-regulated. All fluctuations in the expression levels in the experimental groups were significantly different from those of the control group (p < 0.05) (Table 2). Generally, found target rRNA expression after application of oligonucleotide insecticides corresponds to scheme of DNA containment mechanism (Oberemok and Gal’chinsky 2024; Gal’chinsky et al. 2024). As was seen in our previous experiments, surviving sternorrhynchans after 1^st^ cycle of DNAc mechanism composed of 2 steps (1^st^, ‘arrest’ of target rRNA followed by its hypercompensation and 2^nd^, degradation of target rRNA) tend to overexpress target rRNA (Gal’chinsky et al. 2024) and initiate 2^nd^ cycle of DNAc. Interestingly, 1:1 mixture of Laura-11 and Alacris-11 caused the most pronounced degradation of target rRNA in the pest not allowing to initiate 2^nd^ cycle of DNAc in surviving insects found for other experimental groups on the 14^th^ day (Table 2).

## Discussion

For the first time, this article examines the perspective of using oligonucleotide insecticides and their mixture against psyllids using *T. alacris* as a model insect pest. The results obtained indicate the possibility of development of olinscides against *D. citri* and other economically important representatives of the superfamily Psylloidea. On the one hand, using rRNA as a target makes it possible to generate a powerful insecticidal signal in the cell, and on the other hand, insect rRNA genes have sufficient variability, particularly ITS regions, to create well-tailored species-specific, genus-specific, and family-specific sequences of oligonucleotide insecticides. Of note, oligonucleotide insecticides Alacris-11 and Laura-11, could also be used against other species from superfamily Psylloidea having complete sequence match with them. For example, olinscide Alacris-11 could be successfully used against the olive psyllid *Euphyllura olivina* Costa (family Liviidae) (GenBank sequence ID: MT038970.1; range: 388 to 378). *E. olivina* is considered as one of the most dangerous pests that affect olive trees in the Mediterranean, Palestine (Hamdan and Alkam, 2016) and Taro region of Iran (Khaghaninia, 2009). Also olinscide Alacris-11 could be used against tomato potato psyllid *Bactericera cockerelli* Šulc (family Triozidae) (GenBank sequence ID: MT027594.1; range: 734 to 724). *B. cockerelli* is an important insect pest of potato, tomato, and other solanaceous crops (Sarkar et al. 2023). Olinscide Laura-11 could be used against *Trioza remota*, Foerster (family Triozidae) (GenBank sequence ID: MT038993.1; range: 321 to 311). *T. remota* is a pest which creates galls on the leaves of oak (*Quercus* species) (Redfern et al. 2011). Thus, “genetic zipper” method is not just innovation but also an algorithm which with a high degree of probability calculates the efficiency of a particular olinscide not only for the target insect pest but also for closely related species having perfect complementarity to the developed olinscide.

Why there is a strong confidence that closely related species of target insect pests will be also successfully controlled by oligonucleotide insecticides in the case of perfect complementarity match between a developed DNA olinscide and rRNA of a particular insect pest? Recently, in Frontiers of Agronomy we published opinion article on successful list of pests targeted by oligonucleotide insecticides (Oberemok et al. 2024), and all 13 oligonucleotide insecticides were selected from the first time and did work very efficiently; in average, the developed olinscides caused 80.04 ± 12.73 % mortality for sternorrhynchans in 3-14 days after single or double treatment. With high level of probability all above-mentioned insect pests will be successfully targeted by Alacris-11 and Laura-11 as well. Thus, ‘genetic zipper’ method is a very time-saving approach. Oligonucleotide insecticides is not only new class of insecticides, it is an easy algorithm for fast creation of plethora of species-specific, genus-specific and family-specific pest control agents based on sequences of rRNAs of insect pests.

Chemical insecticides have always made the greatest contribution against insect pests in plant protection. Chemical insecticides are affordable and fast in action because they are small molecules that, having an affinity for a certain component of the cell, disable its functional activity. The main problems connected with the use of chemical insecticides are the emergence of resistance and lack of selectivity in action. Of note, psyllids in native habitats are an important alternative resource for key natural enemies of pest psyllids (Horton, 2024). Oligonucleotide insecticides, due to their unique sequences of nitrogenous bases, are able to provide both, selectivity in action and avoidance of target-site resistance by insect pests (if conservative sequences of target genes are used). The length of an oligonucleotide insecticide ∼ 11 nt makes it possible to create selective oligonucleotide insecticides with a uniqueness frequency equal to 1/4.19·10^6^ (Oberemok et al. 2022). This allows to create well-tailored DNA insecticides for a large group of organisms located in a certain ecosystem at the moment. In the case of ecosystems with increased diversity, such as forests, it is possible to increase the length of oligonucleotide insecticides to 15–20 nt. Taking into consideration such genetic mechanisms as mutations, random assortment of homologous chromosomes during meiosis, crossing over, amplification of genes of cytochrome P450 monooxygenases, etc., it is impossible to imagine how an insect pest can be deprived of the opportunity to generate insecticide resistance (Oberemok et al. 2017). If target-site resistance occurs, new efficient olinscides can be easily re-created displacing target site to the left or to the right from the olinscide-resistance site of the target rRNA (Gal’chinsky et al. 2024; Oberemok et al. 2024).

Oligonucleotide insecticides can be designed using DNAInsector program (dnainsector.com) or manually using sequences of pest rRNAs found in GenBank database. All these characteristics make oligonucleotide insecticides ‘easy insecticides’, and ‘genetic zipper’ method is perceived as an easy-tuned algorithm for plant protection. Oligonucleotide insecticides, as a new class of insecticides, found a convenient developmental plane when we simplified the system to the minimum combination of nitrogenous bases that can cause an insecticidal effect. Using short antisense oligonucleotides of 11–20 bases in length, it is possible to study the basic patterns of action of insecticides based on antisense DNA and further extend this knowledge to more complex options for creation of such pest control agents. A researcher anywhere in the world can independently synthesize the desired olinscide using a DNA synthesizer. Moreover, research teams in each botanical garden or institute dealing with plant protection can synthesize their own arsenal of DNA insecticides that are relevant specifically for their goals. These are new opportunities that did not exist before.

In addition to better target specificity, antisense oligonucleotides are easier and cheaper to synthesize than siRNAs or dsRNAs, and in the human system, antisense oligonucleotides have been shown to have low immunoreactivity, which is also of significant importance for potential applications in edible plants that could accumulate this kind of means of insect pest control. It is important to note, from the point of view of environmental risks, there are no findings in the numerous human clinical studies that prove, for example, genomic integration events attributable to the use of antisense oligonucleotides (Gruber et al. 2023).

It seems that for the first time in the history of plant protection we deal with easy algorithm for creation of insecticides based on rRNA sequences of insect pests. Although oligonucleotide insecticides have proven themselves very well so far only against sternorrhynchans (Hemiptera) and spider mites (Oberemok et al. 2024) more complex formulations of olinscides with auxiliary substances may be effective in controlling pests from other orders of insects.

## Conclusion

During the study, 11nt long olinscides Alacris-11 and Laura-11 showed high efficiency in *T. alacris* control as well as mixture of these olinscides in 1:1 ratio, which demonstrated the highest insect pest mortality and comprised 98.72 ± 1.14% on the 14^th^ day. The selectivity of olinscides was shown by the use of random oligonucleotide CTGA-11, which had no significant insecticidal effect on the bay sucker. Oligonucleotide insecticides acted on *T. alacris* according to DNA containment mechanism in sequence-specific manner. The experimental results show the selectivity and efficiency of olinscides providing a unique platform to create well-tailored oligonucleotide insecticides based on data of rRNA sequences of pests. It is amazing that minimalist approach of ‘genetic zipper’ method, short antisense DNA dissolved in water, is so powerful and eco-friendly innovation against sternorrhynchans, providing new opportunities in plant protection. While today public trust in pest management tools is low, eco-friendly olinscides have a chance to eclipse old tradition and create a new one connected with safety of pest control agents.

## Acknowledgments

We thank our many colleagues, too numerous to name, for the technical advances and lively discussions that prompted us to write this article. We apologize to the many colleagues whose work has not been cited. We are very much indebted to all anonymous reviewers and our colleagues from the lab on DNA technologies, PCR analysis, and creation of DNA insecticides (V.I. Vernadsky Crimean Federal University, Department of Molecular Genetics and Biotechnologies), and OLINSCIDE BIOTECH LLC for valuable comments on our manuscript.

